# Ethnobotanical survey of plant mosquito repellents: Knowledge, utilization, and application methods for malaria prevention in the Rwenzori Region, Western Uganda

**DOI:** 10.64898/2026.05.04.722777

**Authors:** Tony Mugisa, Emmanuel Kimera, Ikiriza Antony, Kakonge Nathan, Kyomuhendo Meble, Maureen Andinda, Charles Idehen, Chinyere Anyanwu, Hussain Yahaya Ungokore, Matthew Chibunna Igwe

## Abstract

**Background:** Malaria remains a major public health challenge in Uganda, particularly in rural areas where access to conventional vector control tools is limited. Communities often use locally available plants as mosquito repellents, but documentation of the specific plants used, their utilization levels, and application methods in the Rwenzori region are limited. This study aimed to identify the types of plants used locally to repel mosquitoes, assess the level of utilization of plant-based mosquito repellents, and determine the methods of application employed by communities.

**Methods:** A community-based cross-sectional study was conducted from June to December 2024 in the seven districts and one city of the Rwenzori region, Western Uganda. Multi-stage sampling was used to select 173 household heads. Data were collected using a pre-tested, translated (Runyoro, Rutooro, Lukonzo) KoboCollect questionnaire and analyzed descriptively with SPSS version 23.

**Results:** Eighty-six percent of respondents reported using plant-based mosquito repellents, with 55% relying exclusively on plants. The most used plants were *Cymbopogon citratus* (citronella/lemon grass, 39.9%), *Rosmarinus officinalis* (rosemary, 25.7%), and *Eucalyptus spp*. (24.3%). The primary application method was planting repellent plants around the house (51.4%), followed by hanging injured plant parts in windows and doorways (28.4%). Other methods included burning or crushing plant parts and applying extracts/oils.

**Conclusion:** Plant-based mosquito repellents are widely used in the Rwenzori region. This study documents community knowledge and practices that could inform future integrated vector management strategies. Further research is needed to evaluate the entomological and epidemiological effectiveness of the plant repellents that are most used plants and the methods commonly applied.

## Introduction

Malaria remains one of the most significant infectious disease challenges globally, with the WHO African Region bearing the greatest burden. According to the World Malaria Report, there were an estimated 263 million malaria cases and approximately 597,000 deaths worldwide in 2023, with a slight increase to 282 million cases and 610,000 deaths reported for 2024 (1;2;4;14).

Uganda is among the 11 African countries that together account for about two-thirds of the global malaria burden, with the entire population of approximately 48 million at risk (2;15;28).

Despite substantial investments in core interventions such as LLINs, IRS, and case management, malaria transmission persists at high levels in many rural areas due to factors including insecticide resistance, outdoor biting behavior of vectors, and challenges in consistent access and use of conventional tools (3;5;6).

In response, communities in malaria-endemic settings often turn to locally available plants with reputed repellent properties as affordable, culturally acceptable supplements or alternatives. Ethnobotanical studies across Africa have documented numerous plant species used traditionally to repel mosquitoes and other insects (27;21). Common examples include species from the genera *Cymbopogon, Eucalyptus, Ocimum, Lantana*, and *Rosmarinus*, which contain essential oils rich in compounds such as citronellal, 1,8-cineole, and linalool (7;11;12;20;28).

In East Africa, including parts of Uganda, Kenya, Ethiopia, and Tanzania, burning, crushing, or planting such species around homes are frequently reported practices (8;9;10).

However, detailed information specific to the Rwenzori region a highland area in Western Uganda characterized by rural livelihoods, diverse ethnic groups, and intense malaria transmission is scarce (13;16;25;26). This study addresses this gap through three specific objectives: [1] identify the types of plants used locally to repel mosquitoes, [2] to assess the level of utilization of plant-based mosquito repellents, and [3] to determine the methods of preparation and application employed by communities in the context of malaria prevention. By documenting these community practices, the study contributes to the broader understanding of indigenous knowledge that may inform future complementary vector control approaches.

## Materials and Methods

### Study design, setting, and population

This was a community-based cross-sectional study conducted in the Rwenzori region of Western Uganda, which comprises seven districts and one city. The region is predominantly rural, with subsistence agriculture as the main livelihood and a documented high burden of malaria. Data collection took place between June and December 2024.

#### Sampling procedure

A multi-stage sampling technique was employed. In the first stage, seven districts were selected by simple random sampling (lottery method) from the nine districts in the region; the city was included purposively to capture urban characteristics. In subsequent stages, sub-counties, villages, and households were selected randomly, with probability proportional to population size where census or village lists were available. At each level, simple random sampling was conducted using a lottery draw (numbered papers placed in a container and drawn blindly).

Within selected households, the household head (aged ≥18 years) was the primary respondent. If unavailable, any other adult resident ≥18 years who was mentally competent and willing to participate was interviewed.

Inclusion criteria were residency in the selected village, age ≥18 years, mental competence, and provision of informed consent.

### Sample size determination

The sample size of 173 was calculated using the Kish-Leslie formula for estimating proportions in a cross-sectional survey:

n = Z^2^ × p × (1 – p) / d^2^ Where Z = 1.96 (for 95% confidence level), p = 0.5 (assumed maximum variability, as specific prior utilization data for this setting were limited), and d = 0.05 (margin of error). A 10% adjustment was applied for potential non-response, and considerations for the multi-stage cluster design (design effect) were incorporated in final adjustments (30).

### Data collection

A structured questionnaire (see S1 Appendix) was developed, pre-tested, and translated into the commonly spoken local languages (Runyoro, Rutooro, and Lukonzo). Trained research assistants administered the questionnaire face-to-face using the KoboCollect mobile application. The tool collected socio-demographic data and information on plant types used, frequency of utilization, reasons for use, preparation methods, and application practices. Data quality checks for completeness and consistency were performed on-site.

#### Data management and statistical analysis

Data were exported from KoboCollect to Microsoft Excel for cleaning and then imported into SPSS version 23 for analysis. Only descriptive statistics were performed (frequencies, percentages, means, and standard deviations), presented in tables and figures. Inferential statistics were not conducted, as the primary aim was description of community practices rather than hypothesis testing or identification of associated factors.

### Data management and analysis

Data were checked for completeness by research assistants before leaving each household. Data were then coded and entered in to the excel spreadsheet. This was then exported SPSS version 23 for analysis. We conducted descriptive statistics and findings are presented as proportions using tables and figures.

#### Ethical consideration

The study proposal, data collection tools, and consent forms were reviewed and approved by the Mbarara University Research and Ethics Committee (Ref: MUST-2024-1365). Administrative clearance was also obtained from the respective district local governments to facilitate engagement with communities during data collection. Written informed consent was obtained from all participants, and data collection was conducted in a quiet, private setting within the respondents’ households. Each participant was reimbursed approximately 6 USD for their time. The study adhered to the principles of the Declaration of Helsinki.

## Results

### Characteristics of respondents

Most respondents, 122 (70%), were aged between 28 and 47 years. Over half, 116 (67.1%), were male, and most, 131 (75.7%), were married. Nearly half, 81 (46.8%), reported an income of less than 300,000 UGX. The highest level of education for most respondents, 83 (48%), was primary school, compared to 43 (24.9%) who had attained tertiary education.

Regarding occupation, the majority, 65 (37.6%), were peasant farmers, while 22 (12.7%) were civil servants. Other characteristics of respondents are shown in table 1 below.

**Table 1:** Baseline characteristics of the respondents (n=173)

| Variables | Category | Frequency (Percentage) |
| --- | --- | --- |
| Age Group of the Respondents | 18 - 27 | 12(7) |
|  | 28 - 37 | 61(35) |
|  | 38 - 47 | 53(31) |
|  | 48 -57 | 25(14) |
|  | 58 & above | 22(13) |
|  | Mean age (SD) | 41 (13.1) |
| Gender of the Respondents | Male | 116(67.1) |
|  | Female | 57(32.9) |
| Marital status | Divorced | 10(5.8) |
|  | Married | 131(75.7) |
|  | Single | 28(16.2) |
|  | Widowed | 4(2.3) |
| Tribe | Muganda | 2(1.2) |
|  | Mukonzo | 39(22.5) |
|  | Munyankole/Mukiga | 59(34.1) |
|  | Munyooro | 3(1.7) |
|  | Mutooto | 55(31.8) |
|  | Mwamba/Mwizi | 11(6.4) |
|  | Others | 4(2.3) |
| Religion of the respondents | Christian | 140(80.9) |
|  | Moslem | 20(11.6) |
|  | Others | 13(7.5) |
| Average monthly Income | Less than 300,000 | 81(46.8) |
|  | 300,001 - 1,000,000 | 72(41.6) |
|  | > 1,000,000 | 20(11.6) |
| Education level | None | 18(10.4) |
|  | Primary | 83(48) |
|  | Secondary | 29(16.8) |
|  | Tertiary | 43(24.9) |
| Occupation of the Respondent | Businessperson | 33(19.1) |
|  | Civil Servant | 22 (12.7) |
|  | Cultural Leader | 4(2.3) |
|  | Peasant Farmer | 65(37.6) |
|  | Political Leader | 3(1.7) |
|  | Religious Leader | 13(7.5) |
|  | Small Scale Farmer | 33(19.1) |

### Utilization of plants for mosquito bite prevention

Utilization of plants for repelling mosquitoes was assessed in comparison with other synthetic repellents sold on the market in the study area. Results in Figure 1 show that majority of the respondents (86%) used plant repellents with 55% using only plant repellents. Only 14.0% used synthetic repellents alone.

**Figure 1:**
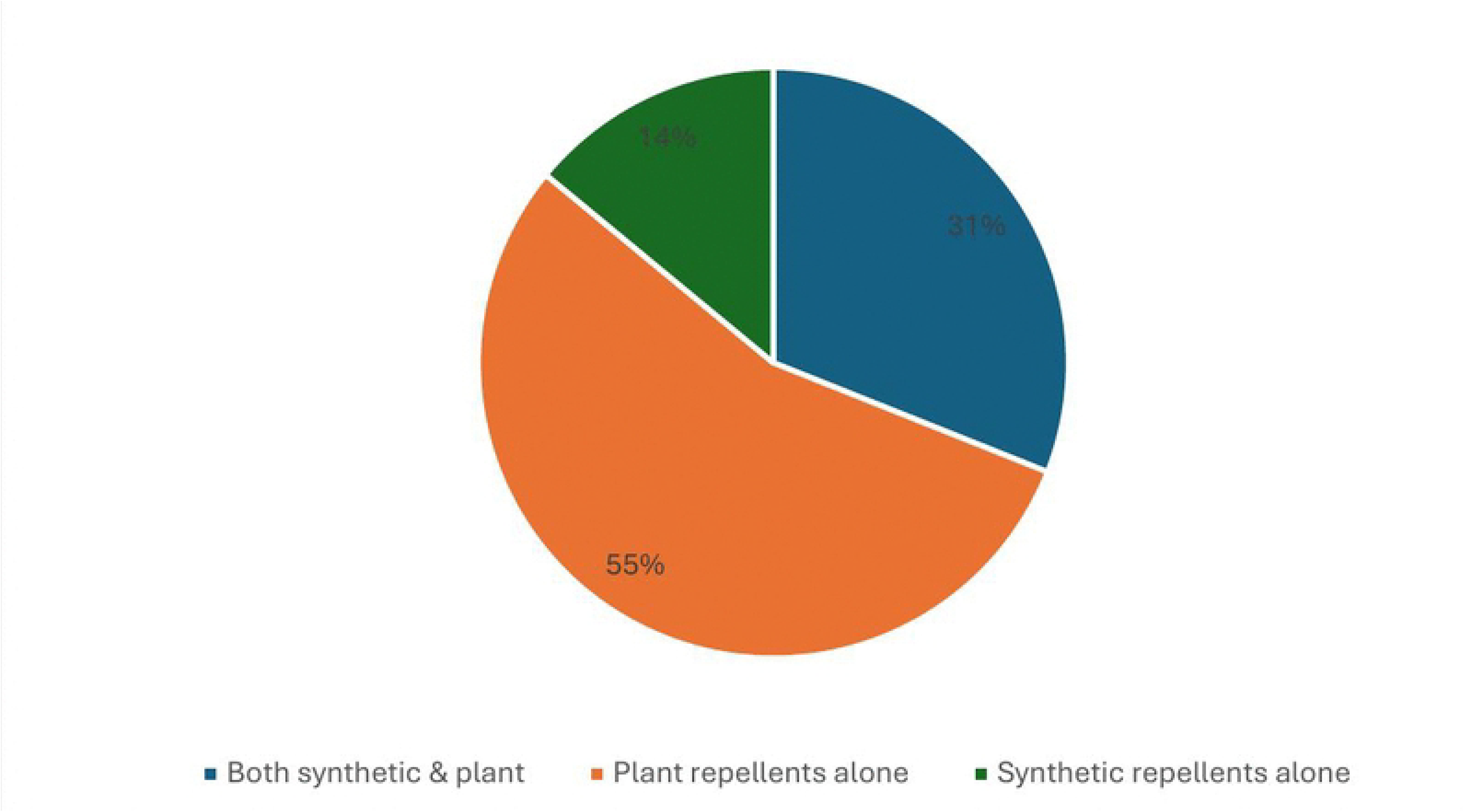
Showing proportion of utilization of plant repellents compared to synthetic repellents in the Rwenzori region (n=173)

### Commonly used mosquito repellant plants

The findings indicate that the most, 59 (39.9%) of the respondents in the study used Citronella (Lemon grass) as their primary mosquito repellent followed by Rosemary 38 (25.7%), Eucalyptus 36(24.3%), *Lantana camara* 32(18.5%), lavender 14(9.5%) Catnip 8(8.1%), Peppermint 32(21.6%), Spearmint 30(20.3%), Marigolds 9(5.2%), *Tetradenia riparia* 12(12.2%), Basil 7(4.7%) while the least 5(7.7%) used Bee balm (Table 2).

**Table 2:**
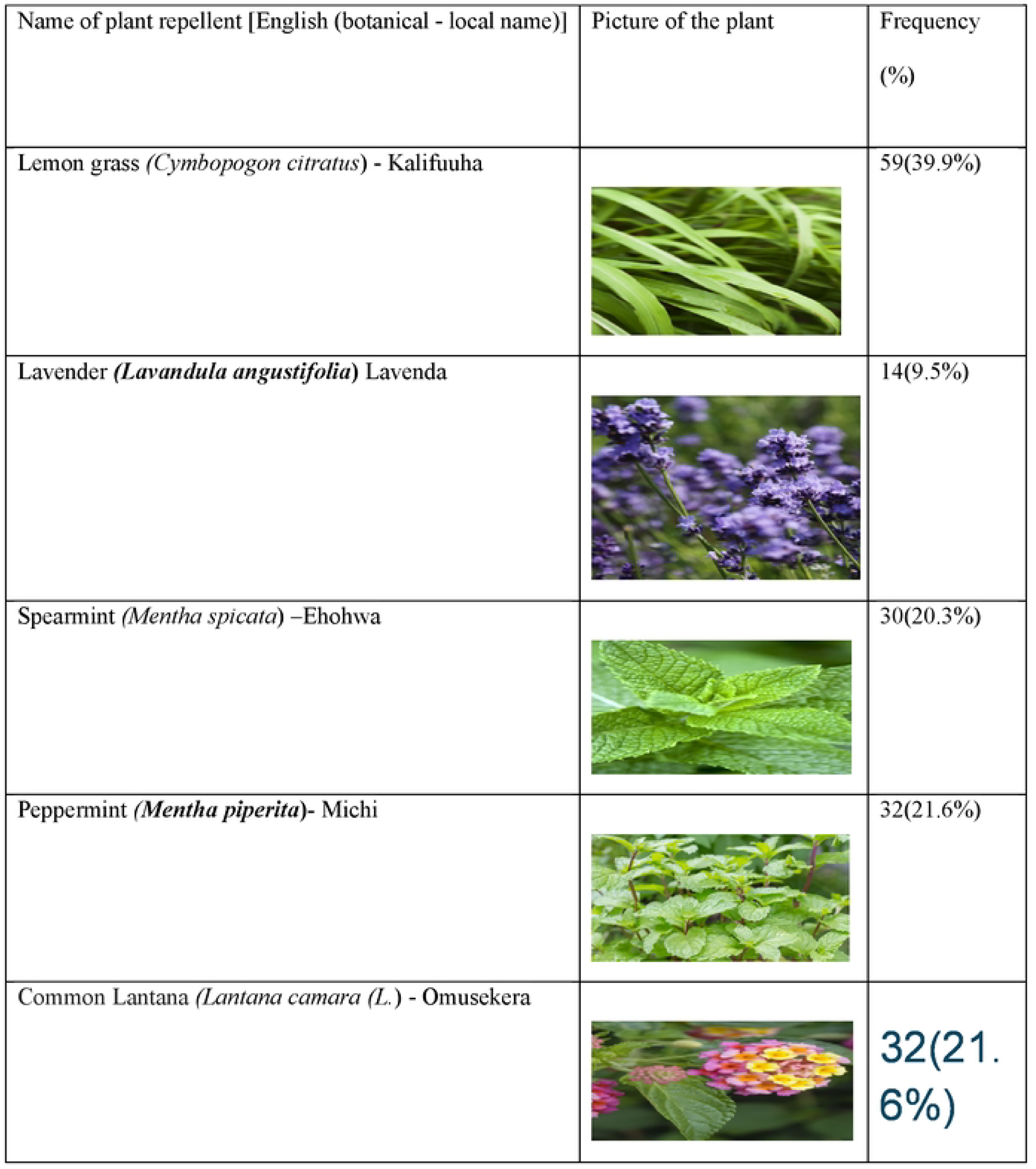
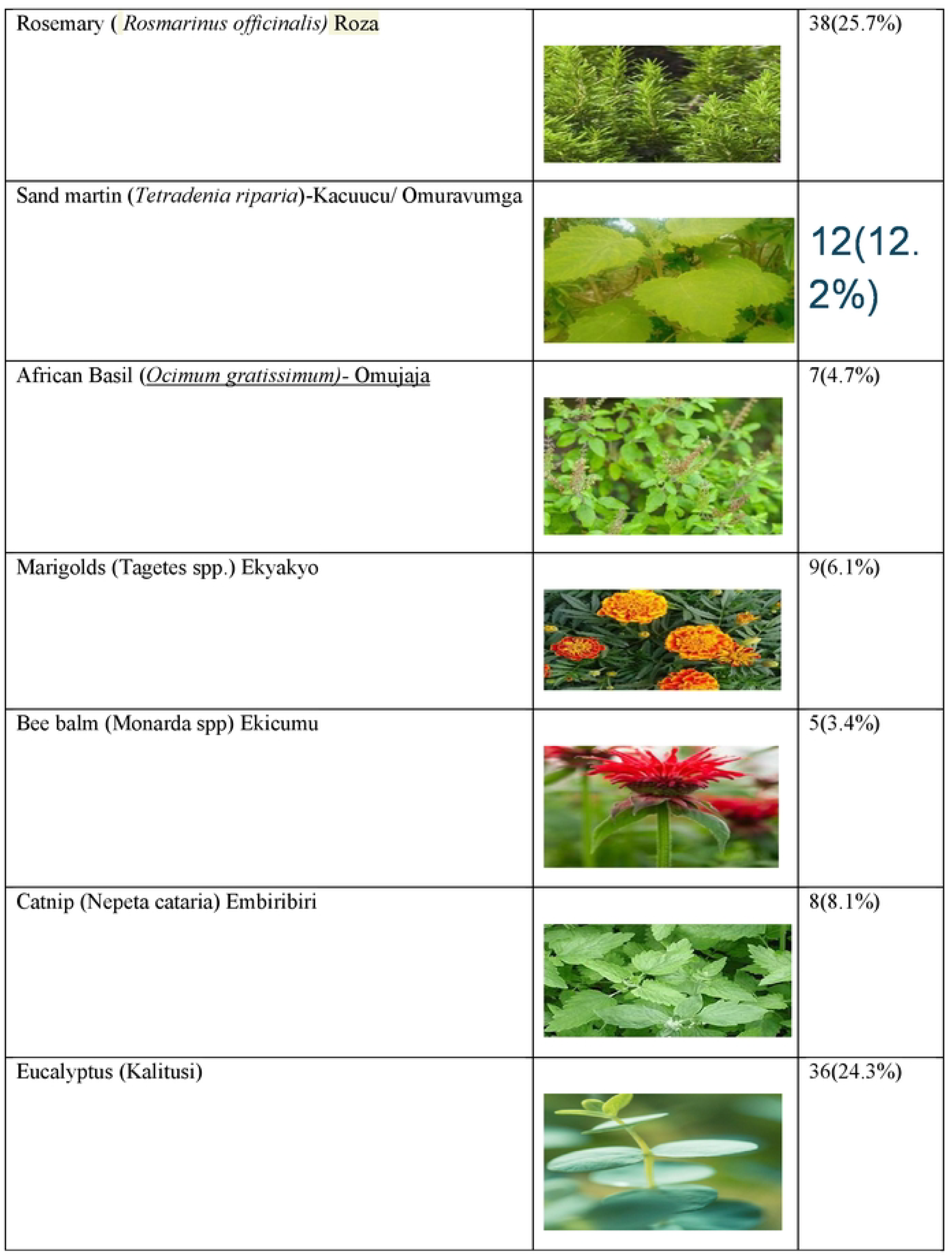
Commonly used plants: Their English names, botanical names, name in local language and reported frequencies.

| Name of plant repellent [English (botanical - local name)] | Picture of the plant | Frequency (%) |
| --- | --- | --- |
| Lemon grass (*Cymbopogon citratus*) - Kalifuuha |  | 59(39.9%) |
| Lavender (*Lavandula angustifolia*) Lavenda |  | 14(9.5%) |
| Spearmint (*Mentha spicata*) –Ehohwa |  | 30(20.3%) |
| Peppermint (*Mentha piperita*)- Michi |  | 32(21.6%) |
| Common Lantana (*Lantana camara* (L.) - Omusekera |  | 32(21.6%) |
| Rosemary ( *Rosmarinus officinalis*) Roza |  | 38(25.7%) |
| Sand martin (*Tetradenia riparia*)-Kacuucu/ Omuravumga |  | 12(12.2%) |
| African Basil (*Ocimum gratissimum*)- Omujaaja |  | 7(4.7%) |
| Marigolds (Tagetes spp.) Ekyakyo |  | 9(6.1%) |
| Bee balm (Monarda spp) Ekicumu |  | 5(3.4%) |
| Catnip (Nepeta cataria) Embiribiri |  | 8(8.1%) |
| Eucalyptus (Kalitusi) |  | 36(24.3%) |

**Table 3:** Specific methods of preparation for the different plant repellents (n=148)

| Plant | Method of preparation |  |  |  |  |  |  |  |
| --- | --- | --- | --- | --- | --- | --- | --- | --- |
|  | Burning<br>in the<br>House | Crushing<br>in the<br>House | Applying<br>Extract/o<br>il on skin | Hanging<br>plants in<br>the doors<br>&<br>windows | Plant<br>around<br>the<br>house | Applying<br>/xtract on<br>surfaces | Candle<br>s<br>enriche<br>ment | Soak in<br>Cotton<br>&hang<br>in doors |
| Basil | 0(0.0) | 0(0.0) | 5(71.4) | 1(14.3) | 1(14.3) | 5(71.4) | 0(0.0) | 0(0.0) |
| Bee Balm | 0(0.0) | 3(60.0) | 0(0.0) | 0(0.0) | 5(100) | 2(40.0) | 0(0.0) | 0(0.0) |
| Citronella | 0(0.0) | 18(30.5) | 35(59.3) | 17(28.8) | 31(52.5) | 33(55.9) | 8(13.6) | 0(0.0) |
| Eucalyptus | 2(5.6) | 0(0.0) | 33(91.7) | 13(36.1) | 2(5.6) | 36(100.0) | 0(0.0) | 0(0.0) |
| Lavender | 1(7.1) | 3(21.4) | 5(35.7) | 0(0.0) | 10(71.4) | 14(100.0) | 0(0.0) | 0(0.0) |
| Marigolds | 0(0.0) | 3(33.3) | 5(55.6) | 2(22.2) | 4(44.4) | 9(100.0) | 0(0.0) | 0(0.0) |
| Peppermint | 5(15.6) | 6(18.8) | 9(28.1) | 0(0.0) | 31(96.9) | 7(21.9) | 0(0.0) | 22(68.8) |
| Spearmint | 0(0.0) | 6(20.0) | 13(43.3) | 4(13.3) | 30(100) | 30(100) | 0(0.0) | 22(73.3) |
| Rosemary | 30(78.9) | 1(2.6) | 0(0.0) | 12(31.6) | 22(57.9) | 24(63.2) | 0(0.0) | 0(0.0) |
| Tetradenia<br>riparia | 0(0.0) | 5(27.8) | 8(44.4) | 0(0.0) | 18(100.0<br>) | 18(100.0) | 0(0.0) | 0(0.0) |
| Catnip | 0(0.0) | 6(50.0) | 12(100) | 0(0.0) | 12(100.0<br>) | 7(58.3) | 0(0.0) | 0(0.0) |
| Lantana<br>Camara | 32(100) | 0(0.0) | 0(0.0) | 0(0.0) | 0(0.0) | 17(53.1) | 0(0.0) | 0(0.0) |
| Total | 70(24) | 51(17.5) | 125(42.8<br>) | 49(16.8) | 166(58.8<br>) | 202(69.2) | 8(2.7) | 22(15.1) |

### Methods of preparation and application by plant species

Findings indicate that the most reported method of preparation for plant repellents is planting the repellent plants around the house 76(51.4.%). Other methods included hanging injured plant parts in the doors and windows of their house 42(28.4%). Additional methods included burning of repellent plant parts in the house 17(11.5%), crushing repellent plant parts in the house 8(5.4%), and application of plant oils 5(3.4%) their bodies and surfaces in the house (Table 2)

## Discussion

This cross-sectional survey documents widespread use of plant-based mosquito repellents among communities in the Rwenzori region, consistent with ethnobotanical findings from other malaria-endemic areas in Africa [9, 11, 12, 18, 25, 26, 27].

The high reported utilization (86%), with over half of users relying exclusively on plants, underscores the importance of traditional knowledge in daily vector avoidance practices where conventional tools may be supplemented or limited by access, cost, or resistance issues. The predominance of *Cymbopogon citratus, Rosmarinus officinalis*, and *Eucalyptus spp*. aligns with global and regional reports highlighting these species for their aromatic essential oils [26, 28, 29].

Planting around homes was the most common method, offering a sustainable, low-cost approach, while other techniques such as hanging, burning, or crushing reflect adaptive local innovations to maximize exposure [19, 20, 29]. These patterns differ somewhat from studies in Ethiopia or Kenya where burning/smoldering was more dominant, possibly due to ecological or cultural variations [10, 11].

## Limitations

Several limitations should be considered when interpreting these findings. The cross-sectional design captures self-reported practices at a single point in time and cannot establish causality or temporal trends. Data are based on self-reports without botanical specimen validation, laboratory confirmation of plant identity, or objective measurement of utilization frequency. Importantly, the study did not assess actual repellent efficacy through entomological (e.g., landing inhibition) or epidemiological endpoints. No control group or direct comparison with synthetic repellents was included. Only descriptive statistics were used; factors associated with utilization were not explored. These constraints mean the results document community knowledge and practices rather than prove effectiveness.

## Conclusion and Recommendations

Plant-based mosquito repellents are widely known and used in the Rwenzori region as part of household malaria prevention strategies. This documentation of commonly used plants, utilization levels, and application methods provides valuable baseline information on indigenous practices. Future studies should prioritize laboratory bioassays, semi-field trials, and standardized efficacy testing (aligned with WHO guidelines) on the most promising species and methods identified here. Such research could support the safe and evidence-based integration of selected plant-based approaches into integrated vector management, complementing existing tools while promoting sustainability and community ownership.

## Declarations

## Abbreviations

ITNS: Insecticide Treated Nets
LLIN: Research and Ethics Committee
Tel: Telephone
LLINS: Long lasting insecticide nets
VHT: Village Health Team
Vs: Versus
WHO: World Health Organization
CI: Confidence Interval
Dr: Doctor
HCIII: Health Center Three
MCD: Malaria Control Division
IRB: Institution Research Board
Km: Kilometers
MoH: Ministry of Health
P-value: Level of Significance
MCP: Malaria control program
PES: Pesticide Evaluation Scheme
NWF: National wildlife federation
IRS: Indoor residual spraying
DEET: N,N-diethyl-meta-toluamide

## Ethics approval and consent to participate

### Norm/Standard according to which research was conducted

This research was conducted adhering to guidelines set forth by the Helsinki Declaration, which outlines ethical principles for medical research involving human subjects.

The study proposal, data collection tools, and consent forms were submitted to the Uganda National Council for research and technology, reviewed and approved by the Mbarara University Research and Ethics Committee (Ref: MUST-2024-1365). Administrative clearance was also obtained from the respective district local governments to facilitate engagement with communities during data collection. Written informed consent was obtained from all participants, and data collection was conducted in a quiet, private setting within the respondents’ households to ensure privacy. Each participant was reimbursed approximately 6 USD for their time.

## Consent for publication

All the authors of this manuscript were fully consulted, and all do consent to publish this article . The study has no identifying images or other personal or clinical details of participants that compromise anonymity. All participants who gave information were consulted and consent to publish the study findings.

### Availability of data and materials

The datasets used and/or analyzed during the current study are available from the corresponding author on reasonable request.

## Author contribution

Tony Mugisa contributed to conceptualization, data curation, analysis, funding acquisition, investigation, project administration, methodology, resources, software, supervision, visualization, and manuscript review and editing. Emmanuel Kimera contributed to conceptualization, data curation, analysis, investigation, methodology, project administration, validation, visualization, and manuscript drafting and review. Mathew Chibuna contributed to conceptualization, analysis, investigation, methodology, validation, visualization, and manuscript drafting and review. Maureen Andinda and Kakonge Nathan contributed to methodology, software, supervision, validation, visualization, and manuscript drafting and review. Antony Ikiriza Chinyere Anyanwu and Hussain Yahaya Ungokore contributed to methodology, software, supervision, validation, visualization, and manuscript drafting and review. Meble Kyomuhendo and Charles Idehen contributed to methodology, supervision, visualization, and manuscript drafting and review.

## Acknowledgement

The Authors thank the research assistants for their integrity and hard work in collecting the data, the Respondents of the nine districts and one city for their cooperation and the Local council leaders provided valuable information for this study. The authors also acknowledge all researchers whose findings helped in the development of this manuscript. The Authors also thank all the study participants-household heads in the nine districts and one city, for accepting to participate in the study.

Appendix one: Questionnaire - English Version

Appendix Two: Informed Consent - English Version

